# Using Disinhibition versus Direct Control in a Spiking Neural Model of Dopamine-Driven Reinforcement Learning

**DOI:** 10.64898/2026.05.22.727086

**Authors:** Roberto Sautto, Nicolas Cuperlier, Thanos Manos, Marwen Belkaid

**Affiliations:** ETIS UMR 8051, CY Cergy Paris Univ., ENSEA, CNRS, F95000 Cergy, France

**Keywords:** Computational Neuroscience, Reinforcement Learning, Spiking Neural Networks

## Abstract

Dopaminergic signalling is central to value learning and decision making. It has been observed that multiple pathways with different patterns of connectivity project to midbrain dopaminergic neurons, some involving direct excitatory projections while others involve disinhibition. However, the respective contributions of these patterns to dopamine control, and their computational and functional advantages remain unclear. In the current work we simulate and evaluate two fully spiking neural models of dopaminergic control, based either solely on disinhibition, or solely on direct inhibitory and excitatory projections. We compare these models in terms of their engineering properties, their resulting spiking profiles, and their ability to successfully acquire representations of expected value in a 3-armed bandit task. We find that both models are able to operate at an asynchronous-irregular firing regime, but that the firing profile of the direct integration model is less resilient to disruption and more sensitive to incoming signals. In addition, the disinhibition model performs better in the learning task. We conclude that while the direct model is more parsimonious, disinhibition-based control remains advantageous in the operational context. Our results have implications for the study of decision-making brain circuits as well as for the design of brain-inspired systems.

## I. Introduction

Making effective decisions in stochastic environments requires forming expectations about the values of available objects and actions based on outcome probability and magnitude. Reinforcement learning has provided a powerful framework to explain the acquisition of such expectations [1]. One of the fundamental notions in this framework is that value representations are updated by calculating so called reward prediction errors (RPE), which are essentially the difference between observed and expected outcomes. There has been tremendous progress in identifying neural correlates of reinforcement learning over the past years. However, it is still unclear how spiking neural networks can implement and use such computations for value coding and decision making.

Dopamine (DA) has been extensively investigated as a neural correlate of reinforcement learning computations. A prominent view holds that dopamine computes RPE, in particular since transient dopaminergic activity was found to carry rich information about magnitude, salience and timing of positive and negative events and outcomes [1, 2]. DA can also affect synaptic strength and thereby drive learning [1, 2]. Moreover, it is worth noting that the tonic dopamine levels have been tied to representations of outcome uncertainty and overall expected reward, and suggested to modulate signal-to-noise ratio and the exploration-exploitation trade-off decisions [3–5]. In general, computational models tend to represent DA activity as a scalar quantity [6–9], or a set firing rate [10]. However, we believe that studying the integration of the neural signals driving dopaminergic activity is paramount to understanding the role of DA-related computations in reinforcement learning.

One key question is what neural architecture is required to implement dopaminergic computations. In other words, assuming that neural populations represent expected and obtained rewards, how should these signals be integrated to produce DA activities enabling reinforcement learning? In the brain, Midbrain dopaminergic neurons (e.g. VTA-DA) are known to be reached by excitatory projections (such as PPTg) [11] on the one hand, and on the other by multiple inhibitory and disinhibitory pathways [11–14] (such as VTAGABA-LHb-RMTg, or NAcc-VP). Similarly, some existing models control DA activity via direct integration of reward and expectation [15], via inhibition-disinhibition [16], or both patterns [17, 18].

Here, we aim to compare the direct and disinhibition-based connectivity patterns and their ability to produce dopamine-driven reinforcement learning. In particular, we ask how these patterns differentially contribute to producing biologically plausible dopaminergic activity [19, 20]. We also examine their potential advantages from the “engineering” perspective, in terms of efficiency, design modularity, and behavioural performance. To test this, we perform two experiments: 1) steady state recording is used to study the firing rates and firing regimes achieved by each model, 2) a probabilistic decision task is simulated to assess the models’ learning abilities. Because disinhibition is less parsimonious, we expect its existence in the brain to be due to functional and computational advantages. Specifically, we hypothesize that:

H1 Disinhibition could better exhibit desired firing rates

H2 Disinhibition could more consistently operate at a desirable firing regime in terms of regularity and synchrony

H3 Disinhibition could better integrate expectation and reward information for successful learning, in terms of distinguishable synaptic encoding of expected values

## II. Model Description

### A. General Architecture and Neuron Model

The general architecture used here builds on previous work modelling a probabilistic decision task with a spiking neural model comprising a decision network modulated by dopaminergic activity driven by preset inputs [21]. Targets with distinct reward probabilities are represented by *c* = 3 population channels per layer, with *n* = 15 neurons each: when a target is available, the corresponding estimation channel receives ∼ 30Hz input. All projections are randomly connected at 70% probability, providing a sufficient amount of synapses for *n* (∼ 157), while avoiding synchrony due to input similarity.

The plastic projection to the estimation layer receives a learning signal from DA and encodes the expected reward, which is reflected in the postsynaptic firing (1-2Hz baseline). A selection mechanism utilising learned values is out of the scope of the present work, and it is substituted by a random choice (50%) between two given options. When *t* is selected the environment layer activates a reward neuron (110Hz) with probability *p*_*t*_ ∈ [25%, 50%, 100%].

The models use LIF neurons (Table I):

**TABLE I.** Parameters at the neuron (top) and network (bottom) levels.

| Neuron Parameters |  | Neuron-type Specific Parameters |  |  |  |
| --- | --- | --- | --- | --- | --- |
| Name | Value | Name | General | Striatal | DA |
| $\tau_{AMPA}$ | 3ms | $E_L$ | -70mV | -80mV | -65mV |
| $E_{AMPA}$ | 0mV | $V_{res}$ | -70mV | -80mV | -65mV |
| $\tau_{NMDA}$ | 100ms | $V_{th}$ | -55mV | -60mV | -50mV |
| $E_{NMDA}$ | 0mV | $\tau$ | 15ms | 10ms | 10ms |
| $\tau_{GABA}$ | 5ms | $R$ | 50MΩ | 60MΩ | 30MΩ |
| $E_{GABA}$ | -70mV | | | | |
| $\sigma$ | 1mV | | | | |

| Shared |  | Disinhibition |  | Direct |  |
| --- | --- | --- | --- | --- | --- |
| Name | Value | Name | Value | Name | Value |
| $I_{Input}^1$ | 0.33 | EST$^-^3$ | 0.4 | EST$^-^3$ | 0.4 |
| $I_{EST}$ | 0.28 | $ID_A^3$ | 0.467 | $ID_A^3$ | 0.412 |
| $I_{Reward}^1$ | 0.8 | Rew-EXC$^3$ | 4.0 | Rew-DA$^3$ | 4.0 |
| Input-EST$^2$ | 0.07 | EST$^+^3$ | 1.0 | EST$^+^3$ | 2.5 |
| $\alpha$ | 0.01 | EXC | 0.24 | | |
| $\tau_{DA}^3$ | 100 | INH | 0.31 | | |
| $\tau_{tonic}^3$ | 500 | EXC-INH | 3.0 | | |
|  |  | INH-DA | 2.0 |  |  |

<sup>1</sup> Only when active.
<sup>2</sup> Initial plastic weight.
<sup>3</sup> Parameters varied in random sampling (see III-D).

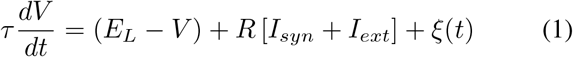

where *V* is the voltage of the neuron, *E*_*L*_ the resting potential, *I*_*ext*_ is the extrinsic current, *τ* is the membrane time constant, and *ξ* is time-resolved white noise with a standard deviation *σ*. A spike is emitted at *V*_*th*_, and *V* is rest to *V*_*res*_. *I*_*syn*_ is the sum of conductance-based synaptic currents *I*_*CH*_ for each ion channel in GABA, AMPA (fast), NMDA (slow):

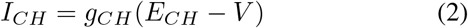

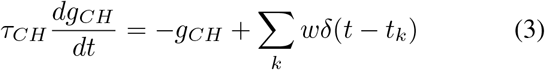

where *g* is a channel’s conductance, *E* is its reversal potential, *τ* is the time constant, *w* is the synaptic strength, and the *δ* function represents the presynaptic spike trains. The slow NMDA channel is always activated by 0.2 times *w* [22].

In order to capture some of the neuronal diversity of the brain circuitry involved in similar functional roles, the model uses three sets of neuron parameters (Table I, top right) based on multiple simulation and recording studies from the literature: the estimation layer is based on striatal and GABAergic neurons, DA on features of VTA/SNc neurons [10, 23], and all other layers use more general parameters.

### B. Neural computation of reward prediction errors

The goal of producing emergent RPE activity in DA neurons and extracting a continuous modulatory signal that conserves both tonic and transient characteristics requires certain features: (a) baseline activity level to capture negative RPE; (b) a threshold to attribute sign to RPE; (c) changes in baseline activity must not interfere with RPE sign.

Summing a positive outcome signal with a negative expectation signal (Fig. 1) shapes DA activity as an uncentered and unsigned RPE. An abstract “DA concentration” can be derived from this activity with a simple moving average maintained by the innervated neuron *i*:

**Fig. 1.**
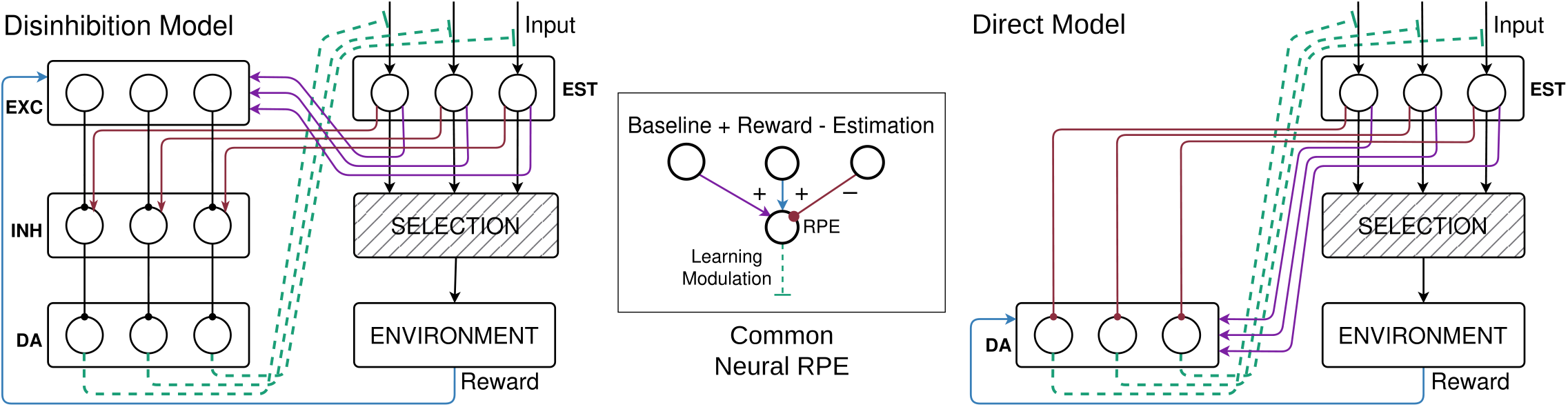
Disinhibition-based and Direct control models. Selection is substituted by a 50% chance choice among two out of three options. Inset: general (colour-coded) illustration of the neural RPE computation implemented in both models. EST: estimation; EXC: excitatory; INH: inhibitory; DA: dopamine

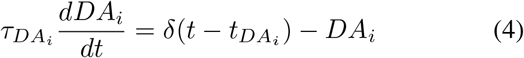

where the *δ* function represents DA spikes that reach *i*.

To attribute a sign to this quantity, we can extract the transient *change* in *DA* by centering it with a threshold [24]. However, DA activity in our models is influenced by contextual information about average expectation, itself dependent on the plastic synaptic strengths. As such, a threshold could lead the system to a feedback loop of constant negative (positive) RPE if the average expectation is low (high).

Our solution is to use tonic DA as a moving threshold to capture ongoing contextual modulation and isolate phasic activity. Tonic DA can be calculated as a long-term average of *DA* (similar to eligibility over timescales in [15]):

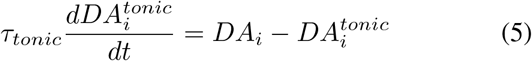

This assumes a homeostatic mechanism, the possible biological details of which are out of scope for this paper, that habituates synapses to DA concentration over time (e.g. availability of postsynaptic messengers), allowing learning to be driven primarily by transient DA changes. Thus, synaptic plasticity is proportional to postsynaptic phasic DA concentration:

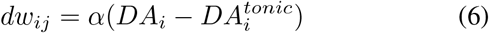

where *α* is a learning rate.

### C. Disinhibition-based and Direct Control Model

The Disinhibition-based control model (Fig. 1, left) utilises two connected inhibitory components to control DA activity, providing a modular mechanism for integrating and regulating multiple contextual and transient signals affecting DA release.

*I*_*EXC*_ and *I*_*INH*_ are tuned for activities at rest of 1Hz and 20Hz respectively, putting DA under constant noisy inhibition to reduce its intrinsic activity (∼ 20Hz) to levels recorded in vivo (∼ 5Hz). The expected reward (Fig. 1, red) excites the inhibitory layer, and DA neurons are further inhibited; reward and average expected reward (Fig. 1, blue and purple) increase excitatory layer activity, which reduces firing in the inhibition layer thus yielding a net depolarising effect on the DA layer.

Average expected reward information is diffused via untargeted projections to estimation, while reward and expected reward information is integrated per-channel. This design abstracts any other eventual credit assignment or selectivity mechanisms that are outside the current scope.

The Direct control model (Fig. 1, right), instead, delivers signals via both excitatory and inhibitory projections: reward and expected reward information is integrated directly at the single-neuron level in the DA layer. As a consequence, the Direct model has fewer degrees of freedom (the excitatory and inhibitory layers need four additional parameters) and further detailed modulation of excitatory and inhibitory influences is left to the areas that generate them. Similarly, the baseline firing rate of DA neurons is mainly controlled by the extrinsic current and average expected reward.

## III. Methods

### A. Implementation Implementationand Simulation

The models were implemented in Python with the BRIAN2 [25] neural simulator, using the C++ code-generation feature for high computational efficiency.^1^

We used ad-hoc abstractions of putative signal gating mechanisms for reward and expectation signals: the corresponding projections, as well as plasticity, are only active for a certain period of time after selection, and only for the selected channel. This was achieved by exploiting the event-trigger nature of neuron models to set flags as well as to activate/deactivate the desired input populations at the start of each trial.

### B. Experiment 1: Steady State Recording

We turned off the selection and plasticity mechanisms, and recorded the spike trains of the DA neurons with all inputs active. Each 10s recording was preceded by 10s habituation to allow slow currents to settle in their regimes.

Behaviours under different operational conditions were examined by varying expected reward (input-estimation synaptic strength: low = 0.04, high = 0.1) as well as the presence of reward (no: *I*_*Reward*_ = 0, yes: *I*_*Reward*_ = 0.8) for all channels.Each condition was simulated for *s* = 30 sessions. From the resulting recordings we computed various metrics at a channel level, yielding *s* × *c* = 90 data points per condition:

#### Firing Rate

The spike count was divided by the recording duration for each neuron, and then averaged per channel. DA is expected to show resting rates around 5Hz (modulated by average expected reward), and increase with reward [19].

#### Coefficient of Variation

CV measures firing regularity [26] as CV = *σ*_*ISI*_*/μ*_*ISI*_, where *σ* is the standard deviation and *μ* is the mean of the interspike interval (ISI) histogram of a given neuron. A CV = 1 indicates a poisson-like irregular firing; lower values (CV *<* 0.8) indicate a regular firing regime, while higher values (CV *>* 1.2) indicate highly irregular or bursty firing. CVs were calculated per neuron and averaged per channel. DA neurons are expected to show irregular firing that doesn’t necessarily regularise in rewarding conditions [19] if not within bursts (not modelled in the present work).

#### SPIKE-distance

This is a time-resolved synchrony metric using dissimilarities in spike times at each time step [27]. It was calculated at channel level with the library PySpike [28]. Without specific reference points, this is used as a comparative measure of time-based synchrony between the two models.

#### Population Synchrony

In [26] spike count histograms are used to measure multiunit synchrony as 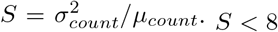 indicates an asynchronous population. Due to the lower population size and firing rates, we could not use the 3ms bin size, opting for multiple bin sizes in 20ms steps up to 100ms. In general, DA neurons are expected to be asynchronous and increase their synchrony during rewarding events [20].

### C. Experiment 2: Learning Task

We simulated a three-armed bandit task [21] by sequentially activating two of the three channels to present all pairs of targets evenly. Since it is not the focus of this paper, action selection was substituted by a random choice (Fig. 1). Selection is simulated after 1s of activity and learning is turned on for 200ms, and a resting period of 500ms with inactive inputs concludes the trial. *t* = 90 trials are run for each of the *s* = 30 sessions and the final synaptic weights of the input-estimation projection are averaged by channel.

Assuming a direct relationship between acquired expectations and action selection, the slope of the linear fit of the weights with the reward probability of the targets is an indicator of exploitative behaviour. For this relationship to be meaningful an action selection system needs to receive discriminable information about expected reward: as such we define learning to be “successful” if the weights of the channels are pairwise significantly different.

### D. Parameter Space Sampling

The base parameters for the two models (in Table I, bottom) are tuned to result in weights around 0.03 for the *p* = 25% target and around 0.10 for the one with *p* = 100%. We tested the resilience of each model, meaning the extent to which their behaviour is consistent (architectural effects) or variable (parameters effects), by varying model parameters that directly affect signal integration and learning behaviour (Table I, 3). Furthermore, exploring the parameter space allows us to show the ability of the models to support targeted disruption (e.g. lesions, chemical blockades).

For these reasons, both Experiment 1 and 2 are repeated with multiple variations (*v* = 100). In hyperparameter optimization, it has been shown that random search is a more efficient way to cover the parameter space than a classical grid search [29]. Here, we use a similar approach: for each variation, each of 6 parameters is sampled from a range of ±10% of their base value.

## IV. Results

### A. Experiment 1

Fig. 2 shows an overview of experimental results. An OLS linear model *model* × (*reward*+*expected reward*) (baseline: Direct, no reward, low expected reward) was fit in order to properly compare behavioural effects of the four conditions: all results reported below are significant (*p <* 0.001). When normality cannot be guaranteed metrics are reported as mean/median unless otherwise stated.

**Fig. 2.**
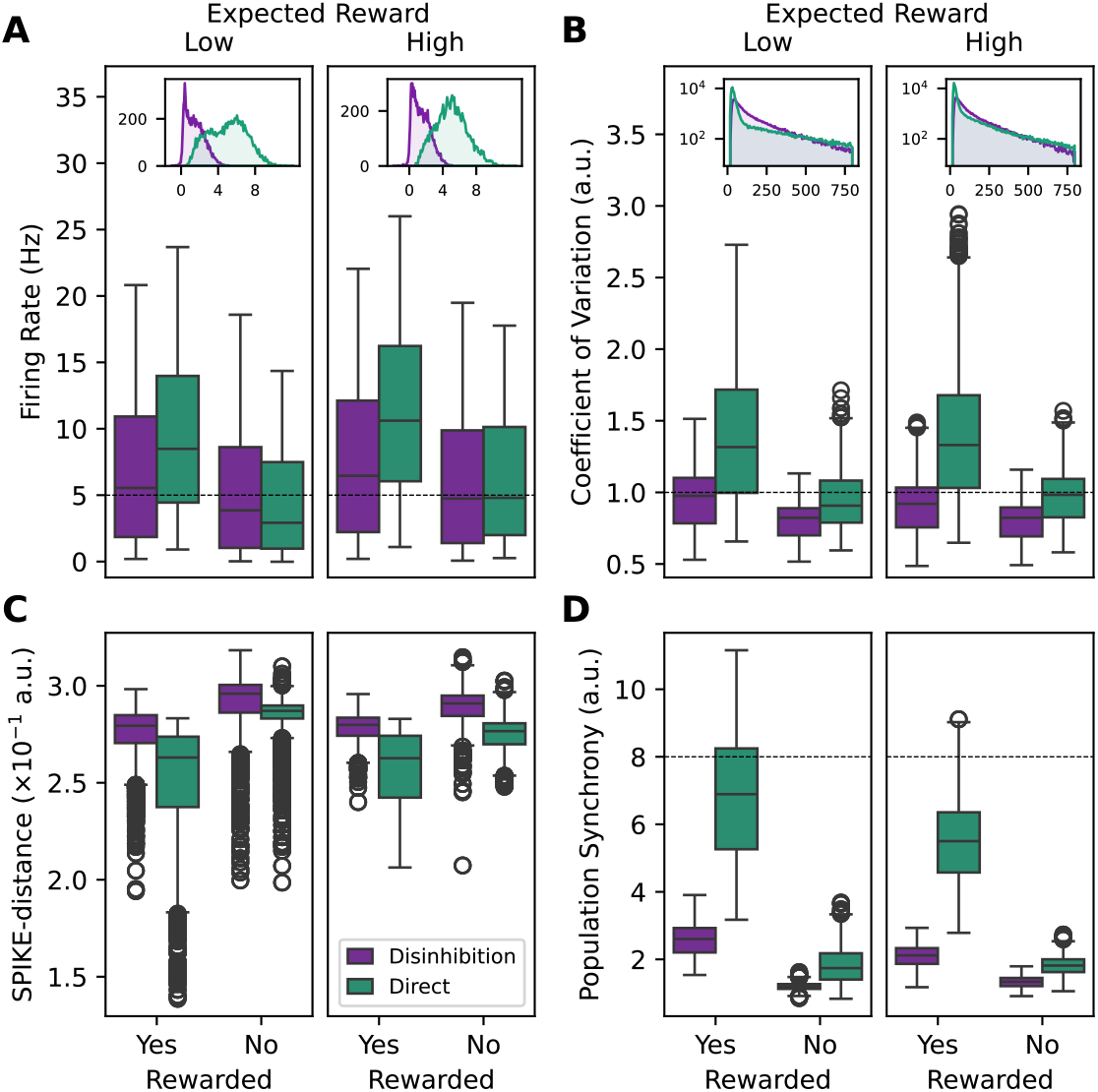
Firing activity metrics for Experiment 1 under the four conditions over parameter variations. **A**: average firing rate; in the inset a distribution of the difference in rate between the rewarded and unrewarded conditions. **B**: Coefficient of Variation (irregularity); in the inset a log-scale distribution of ISIs for the rewarded condition. **C**: SPIKE-distance (asynchrony). **D**: Population Synchrony as in [26], with 20ms bins. Each boxplot shows: median (horizontal line), IQR=Q3-Q1 (box).

#### Firing Rate

Both models are able to produce resting activities (Fig. 2A) within the acceptable range (Disinhibition: 5.85/4.34Hz; Direct: 5.42/4.17Hz). Firing rates increase with reward (*β* = 5.01, *R*^2^ = 0.134) but the effect is severely lessened for the Disinhibition model (*β* = − 3.52). Similarly, both models are being modulated by the level of expected reward (*β* = 1.85), but the effect is weaker in the Disinhibition model (*β* =− 1.08). A distribution of the difference in firing rate between the rewarded and unrewarded condition (Fig. 2A, inset) reveal a peak around 0Hz for the Disinhibition model, and around 4Hz for the Direct model. This suggests that disruption in the Base parameters causes reward integration failure in the Disinhibition, but not in the Direct model, partially explaining the lower sensitivity to reward.

#### Coefficient of Variation

Median firing variability (Fig. 2B) is in the range of an irregular firing regime (0.8-1.2) for both models at rest (Disinhibition: 0.82; Direct: 0.944), but reward injection causes the Direct model to reach high irregularity (1.38/1.33), which can still be acceptable, but is not predicted here without an explicit model of bursting behaviour. The OLS model (*R*^2^ = 0.396) reveals that the average expected reward slightly increases irregularity in the Direct model (*β* = 0.017) but the effect is reversed by the Disinhibition model (*β* = − 0.043), indicating that the filtering of this baseline signal in the latter has a slight regularising effect on the resulting activity. The insets in Fig. 2B corroborate this, showing a peak for short latencies in the log scale ISI distribution of the Direct model, which is much less pronounced or absent in the Disinhibition model, meaning the latter is closer to an exponential ISI distribution and thus poisson-like irregularity.

#### SPIKE-distance

Temporal dissimilarity as measured by this metric (Fig. 2C, *R*^2^ = 0.426) indicates that activity in the Disinhibition is more asynchronous (*β* = 0.01), and that the expected synchronising effect of reward is present in the Direct model (*β* = − 0.025) but is weaker in the Disinhibition model (*β* = 0.011). A very weak effect is seen for expectation (*β* = 0.002) but is absent in the Disinhibition model.

#### Population Synchrony

As seen in Fig. 2C (*R*^2^ = 0.844), population synchrony measured with 20ms bins agrees with SPIKE-distance metrics: the Disinhibition model is more asynchronous overall (*β* = − 0.79); synchrony increases with reward strongly for the Direct (*β* = 4.35) and more weakly for the Disinhibition model (*β* = − 3.27). The average expectation level has a considerable synchronising effect (Direct: *β* = −0.65, Disinhibition: *β* = 0.47). Interestingly, both models remain in the asynchronous regime (as defined by [26]) even with reward, but for any other bin size *>* 20ms (not shown) the metric is ≫ 8 for the rewarded Direct model, while other conditions never are.

### B. Experiment 2

Fig. 3 (A, B) shows the results of the Learning Task for all parameter sets samples. As it is apparent, the weights learnt by the Disinhibition model remain in a more restricted range than those of the Direct model, resulting in a more consistent operational range even after disruption. At the contrary, slope distribution (Fig. 3A and B, inset) for the first model span a larger range than the second, which clusters around 0.08.

**Fig. 3.**
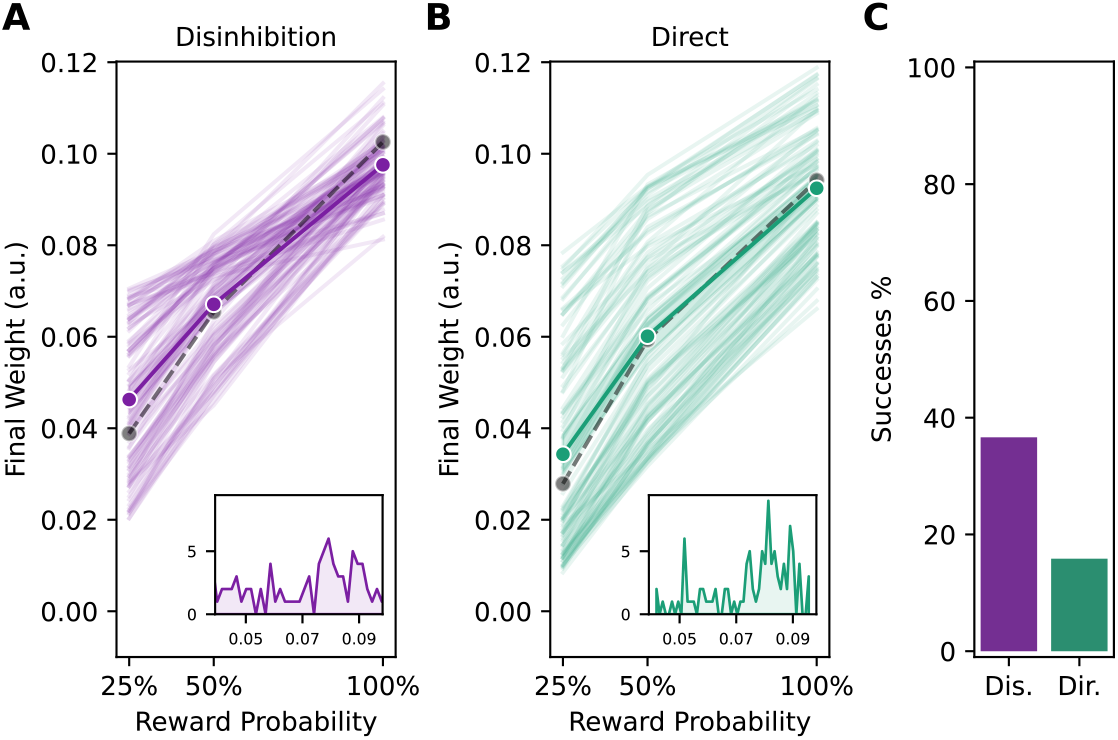
Metrics for the Learning Task over the parameter space samples. **A, B**: relationship between average weights after learning and the reward probability of the target; coloured dots as the average over parameter samples, gray dashed line showing the base parameters; in the inset, distributions of the slopes. **C**: percentage of the parameter samples that resulted in successful learning.

37% of the parameter sets produced successful learning (pairwise significantly distinct weights) in the Disinhibition model, while only 16% were successful in the Direct model (Fig.. 3C). Furthermore, the failed parameter sets of the Direct model contain 77% of all the significantly distinct weight pairs considered, while those failed in the Disinhibition model contain only 50%, suggesting that the latter shows the desirable functional property of failing hard, while the former can fail silently or only partially: this is especially important when considering cases where the relationship between synaptic weights representing expected reward and choice behaviour are direct but not linear (e.g. modulation of action selection), as distinguish from failures in learning.

## V. Discussion

In this paper, we sought to compare two architecturally distinct spiking neural models respectively implementing a disinhibition-based and direct control of DA activity. Our goal was to assess these two models in terms of biological plausibility as well as engineering properties. Assuming the less parsimonious disinhibition pattern brings about computational and functional advantages, we formulated three hypotheses that we tested through steady state in-silico recordings and a probabilistic reward learning task.

First, results from the steady state recording experiment suggest that the effect of the disinhibition architecture is to dampen incoming signals and reduce the model’s sensitivity while still allowing for appropriate integration. By contrast the direct integration architecture was much more easily swayed by incoming signals. Yet, the Direct model was able to maintain appropriate firing rates at resting and rewarded states, thus disproving H1. Second, both models remained in desirable asynchronous-irregular firing regimes. However, the firing metrics for the Direct model showed a much wider dispersion, while the Disinhibition model retained a more conservative profile over parameter sampling. These results thus provide support for H2. Third, one might assume that a lower sensitivity to reward and expectation information would lead to impaired learning, but this proved not to be the case. Indeed, parameter sampling over the Disinhibition model proved more than twice as successful in learning distinct weights than the Direct model, which confirmed H3.

This work provides insights into computational and functional advantages of disinhibition and open various perspective for future research. First, the higher resilience and hard failures of the Disinhibition model might be related to the conservation of the firing regime. It could also be that it is harder for the Direct model to significantly distinguish weights due to the higher variability in produced activity and learned weights.

Further analysis of weight dispersion, and on the responsibility of the single parameters in producing successful learning needs to be conducted in order to distinguish the two explanations, and in particular to study the effects of the conditions in action selection. Furthermore, the highly irregular firing regime reached by the Direct model appears of particular interest. The possibility that a dynamic, rather than static, interplay between excitatory (with fast and slow component) and inhibitory drive could result in emergent burst-like behaviour warrants more attention, and thus future work might explore a hybrid model where contextual baseline modulation is integrated via direct excitatory projections, or redundant signals are integrated in both architectural modes.

Last, our results might also have implications beyond the study of dopamine control. Indeed, disinhibition is a connectivity pattern found in various areas, including the basal ganglia for example. Understanding whether some of our findings apply in other contexts would provide valuable insights into canonical computations in the brain. It is also for interest from the engineering perspective to drive design choices of brain-inspired systems. The main engineering advantage of the Direct control model is its parsimony both in terms of degrees of freedom (parameters) and moving parts (number of neurons), leading to a model that is intrinsically both easier to optimise and more energy efficient in a neuromorphic implementation. On the other hand, the Disinhibition-based control model is designed as a modular and extendable architecture of multilevel integration of incoming signals, which can prove functionally advantageous if the signals need to be differentially modulated.

In conclusion, controlling DA neuron firing via disinhibition, while computationally more costly and less parsimonious, provides distinct design and functional advantages that make it an architecture preferable to integrate into a wider action selection and neuromodulation model of decision making.

## Acknowledgments

The authors would like to thank Aëlien Moubêche for the precursory work on this project.

## Footnotes

1 Codebase available at github.com/BelkaidLab/sautto2026using_IJCNN

## References

[1] W. Schultz, P. Dayan, and P. R. Montague, “A Neural Substrate of Prediction and Reward,” Science, vol. 275, no. 5306, Mar. 1997.

[2] E. O. Neftci and B. B. Averbeck, “Reinforcement learning in artificial and biological systems,” Nature Machine Intelligence, vol. 1, no. 3, 2019.

[3] J. Naudé, S. Tolu, M. L. Dongelmans, N. Torquet, S. Valverde, G. Rodriguez, S. Pons, U. Maskos, A. Mourot, F. Marti, and P. Faure, “Nicotinic receptors in the ventral tegmental area promote uncertainty-seeking,” Nature Neuroscience, vol. 19, Jan. 2016.

[4] M. C. Avery and J. L. Krichmar, “Neuromodulatory Systems and Their Interactions: A Review of Models, Theories, and Experiments,” Frontiers in Neural Circuits, vol. 11, 2017.

[5] E. Lee, M. Seo, O. Dal Monte, and B. B. Averbeck, “Injection of a dopamine type 2 receptor antagonist into the dorsal striatum disrupts choices driven by previous outcomes, but not perceptual inference,” Journal of Neuroscience, vol. 35, no. 16, Apr. 2015.

[6] A. G. E. Collins and M. J. Frank, “Opponent actor learning (OpAL): Modeling interactive effects of striatal dopamine on reinforcement learning and choice incentive.” Psychological Review, vol. 121, no. 3, 2014.

[7] T. Gilbertson and D. Steele, “Tonic dopamine, uncertainty and basal ganglia action selection,” Neuroscience, vol. 466, 2021.

[8] S. M. Vogt and U. G. Hofmann, “Neuromodulation of STDP through short-term changes in firing causality,” Cognitive Neurodynamics, vol. 6, no. 4, 2012-08-01.

[9] H. Wei, Y. Bu, and Z. Zhu, “Robotic arm controlling based on a spiking neural circuit and synaptic plasticity,” Biomedical Signal Processing and Control, vol. 55, Jan. 2020.

[10] A. González-Redondo, J. Garrido, F. Naveros Arrabal, J. Hellgren Kotaleski, S. Grillner, and E. Ros, “Reinforcement learning in a spiking neural model of striatum plasticity,” Neurocomputing, vol. 548, 2023.

[11] A. A. Grace, S. B. Floresco, Y. Goto, and D. J. Lodge, “Regulation of firing of dopaminergic neurons and control of goal-directed behaviors,” Trends in Neurosciences, vol. 30, no. 5, 2007.

[12] K. Morita, M. Morishima, K. Sakai, and Y. Kawaguchi, “Reinforcement learning: computing the temporal difference of values via distinct corticostriatal pathways,” Trends in neurosciences, vol. 35, no. 8, pp. 457–467, 2012.

[13] S. C. Gantz, C. P. Ford, H. Morikawa, and J. T. Williams, “The Evolving Understanding of Dopamine Neurons in the Substantia Nigra and Ventral Tegmental Area,” Annual Review of Physiology, vol. 80, no. 1, 2018.

[14] P. M. Baker, T. Jhou, B. Li, M. Matsumoto, S. J. Y. Mizumori, M. Stephenson-Jones, and A. Vicentic, “The Lateral Habenula Circuitry: Reward Processing and Cognitive Control,” The Journal of Neuroscience: The Official Journal of the Society for Neuroscience, vol. 36, no. 45, 2016.

[15] P. Berthet, M. Lindahl, P. J. Tully, J. Hellgren-Kotaleski, and A. Lansner, “Functional Relevance of Different Basal Ganglia Pathways Investigated in a Spiking Model with Reward Dependent Plasticity,” Frontiers in Neural Circuits, vol. 10, 2016.

[16] W. Potjans, M. Diesmann, and A. Morrison, “An Imperfect Dopaminergic Error Signal Can Drive Temporal-Difference Learning,” PLOS Computational Biology, vol. 7, no. 5, 2011.

[17] P. Kaushik, J. Naudé, S. B. Raju, and F. Alexandre, “A VTA GABAergic computational model of dissociated reward prediction error computation in classical conditioning,” Neurobiology of Learning and Memory, vol. 193, 2022.

[18] J. B. Inglis, V. V. Valentin, and F. G. Ashby, “Modulation of Dopamine for Adaptive Learning: A Neurocomputational Model,” Computational Brain & Behavior, vol. 4, no. 1, 2021.

[19] M. Marinelli and J. E. McCutcheon, “Heterogeneity of dopamine neuron activity across traits and states,” Neuroscience, vol. 282, pp. 176–197, 2014.

[20] C. Paladini and J. Roeper, “Generating bursts (and pauses) in the dopamine midbrain neurons,” Neuroscience, vol. 282, pp. 109–121, 2014.

[21] M. Belkaid and J. L. Krichmar, “Modeling uncertainty-seeking behavior mediated by cholinergic influence on dopamine,” Neural Networks, vol. 125, May 2020.

[22] A. J. Watt, M. C. Van Rossum, K. M. MacLeod, S. B. Nelson, and G. G. Turrigiano, “Activity Coregulates Quantal AMPA and NMDA Currents at Neocortical Synapses,” Neuron, vol. 26, no. 3, Jun. 2000.

[23] A. A. Grace and S.-P. Onn, “Morphology and electrophysiological properties of immunocytochemically identified rat dopamine neurons recorded in vitro,” Journal of Neuroscience, vol. 9, no. 10, pp. 3463– 3481, 1989.

[24] Q. Zhu, F. Han, Y. Yuan, and L. Shen, “A TAN-dopamine interaction mechanism based computational model of basal ganglia in action selection,” Cognitive Neurodynamics, vol. 18, no. 5, 2024.

[25] M. Stimberg, R. Brette, and D. F. Goodman, “Brian 2, an intuitive and efficient neural simulator,” elife, vol. 8, . e47314, 2019.

[26] T. C. Potjans and M. Diesmann, “The cell-type specific cortical micro-circuit: Relating structure and activity in a full-scale spiking network model,” Cerebral Cortex (New York, N.Y.: 1991), vol. 24, no. 3, 2014.

[27] T. Kreuz, D. Chicharro, C. Houghton, R. G. Andrzejak, and F. Mormann, “Monitoring spike train synchrony,” Journal of neurophysiology, vol. 109, no. 5, pp. 1457–1472, 2013.

[28] M. Mulansky and T. Kreuz, “Pyspike—a python library for analyzing spike train synchrony,” SoftwareX, vol. 5, pp. 183–189, 2016.

[29] J. Bergstra and Y. Bengio, “Random search for hyper-parameter optimization,” The journal of machine learning research, vol. 13, no. 1, pp. 281–305, 2012.

